# Identification of potential distribution area for the sweet cherry (*Prunus avium*) by the MaxEnt model

**DOI:** 10.1101/2023.10.26.564153

**Authors:** Hongqun Li, Xiaolong Peng, Peng Jiang, Ligang Xing

**Author notes:** Corresponding Author Hongqun Li, School of modern agriculture and Bioengineering, Yangtze Normal University, Chongqing 408100, China.

## Abstract

The potential distribution of species offers important information for species management, especially for fruit trees with economic value. The sweet cherry is a tree species with higher economic value among deciduous fruit trees in north China. Herein, its potential distributions were modeled under current conditions. The performances of this model were excellent with AUCs of >0.9 for model training and testing. The key factors identified were the thermal factors (min temperature of coldest month (bio06) from -14.5 to 4.5°C, the mean temperature of warmest quarter (bio10) from 21.0 to 28.0°C), followed by water factor (the annual precipitation (bio12) from 500 to 1200 mm). Its suitable region in China mainly is located on 6 geographical regions including southwest China (eastern Sichuan, northeast and main urban areas of Chongqing, mid-western Guizhou and mid-northern Yunnan), northwest China (mid-southern Shaanxi, southern Ningxia mid-southern and eastern Gansu), northeast China (Coastal region of Liaoning), central China (most of Henan, mid-northern Hubei and central Hunan), north China (Beijing, Tianjing, mid-southern Shanxi), east China (Shanghai, Jiangsu, Shandong, central Zhejiang, central and northern Anhui and eastern Jiangxi) and south China (western Guangxi). Relative to their actual distributions, Hubei, Hunan and Anhui etc have been identified to have some potentially suitable areas for these sweet cherries. Among them, these 9 provinces or cities (Shaanxi, Beijing, Tianjing, Shanxi, Hebei, Henan, Shanghai, Jiangsu and Shandong) are main regions for its current development and utilization, followed by Sichuan, Guizhou,Yunnan, Gansu, Liaoning, Hubei, Zhejiang and Guangxi etc. The results can be adopted to identify the suitable areas for its introduction and cultivation to avoid the losses of human labor, material and financial resources due to improper decision, and to provide a scientific basis for its current introduction, cultivation and management.

## 1. INTRODUCTION

The sweet cherry (*Prunus avium* Linn), belonging to genus Prunus in the family Rosaceae, is one of the earliest mature deciduous fruit trees in northern China, and its fruit ripens early and be rich in nutrients with excellent color, shape, and taste, hence it attracts domestic and international attention from consumer and be called as the “first branch of spring fruit” and the “treasure” in the fruit (Nie *et al*., 2022; Luo *et al*., 2022). Furthermore, it has still high medicinal value for humans due to its abundance of bioactive substances such as vitamin C and polyphenols (Wu *et al*., 2018). In China, the sweet cherry originating from southeastern Europe and west Asia, was introduced to China in 1871 and then became tree species with higher economic benefits, showing a vigorous development trend. For instance, in 2016 its total planting area of China ranking first in the world, has exceeded 0.18 million (hm^2^) and its annual output is about 0.7 million ton (Hu *et al*., 2011; Zhang *et al*., 2017). According to the current planting situation of sweet cherry in China, there are two advantageous cultivation areas for this species such as the Bohai Bay Rim area and the area along the Longhai railway (Sun *et al*., 2016; Zhang *et al*., 2017). For example, some planting areas are found in Shandong (Yantai and Taian), Liaoning (Dalian), Beijing, Hebei (Qinhuangdao), Henan (Zhengzhou), Shaanxi (Xi’an), and Gansu (Tianshui) (Zhang *et al*., 2017). After China’s reform and opening up, rural productivity has been greatly liberated and many people have engaged in efficient production through multiple channels. Especially after the 1980s and 1990s, due to the high yield and high nutritional value of the sweet cherries, they received widespread attention in the cultivation industry. At present, the improvement in people’s living standards, going with development of the global economy, will inevitably lead to increasing demand for the sweet cherry. In order to satisfy the increasing demand of this fruit in China, it is highly necessary to expand the growing region of this fruit (Li & Wang, 2009). However, without scientific planning and professional guidance, some farmers possibly planted it in non optimal growth areas, resulting in a series of problem, such as poor plant growth and relatively low efficiency. To avoid investment risks mainly evoked by blindly expanding its introduction and cultivation, it is extremely important for governments at all levels to carry out the research on the national planting division of the sweet cherry, put forward the index system of its planting division adapted to the climate characteristics of China and form opinions for the zoning of the sweet cherries in China.

Fortunately, with the comprehensive application of statistical tools and geographic information system (GIS), many species distribution models (SDMs) have become the important tools available free for the simulation and visualization of the spatial distribution of the species and also determined key environmental factors that limit a species’ geographical distribution (Guisan & Thuiller, 2005; Wang *et al*., 2017). Specifically, based on some specific algorithm, these SDMs can assess the ecological niche of species by using the objective population geographical location and environmental variables affecting its distribution, and projects it into the environment to reflect the preference of species to the habitat in the form of probability (Zhuang *et al*., 2018; Guo *et al*., 2020). However, out of various SDMs, Maxent model (Maximum entropy model) has been confirmed to outperform other SDMs in predicting accuracy, especially in the case of lacking species occurrence data (Wang *et al*., 2010; Qin *et al*., 2017; Liao *et al*., 2017). The several advantages of this Maxent model are as follows: (1) This model only needs the presence-only occurrence points (no absence data are needed) and environmental variables; (2) This model may use continuous and categorical variables, and overcome the correlations to avoid interactions between variables; (3) The models can identify some mainly environmental factors that are responsible for a species’ distribution; (4) Many studies have also confirmed that the model also has good prediction results in the situation of incomplete data and small sample sizes (Wang *et al*., 2017; Qin *et al*., 2017). Additionally, the demands for computer configuration are relatively low, and it has user-friendly operation interface. Therefore, this model has been extensively adopted in the protection of wild animals and plants, management of Invasive species, transmission of forest diseases and pests, and the habitat suitability analysis of medicinal plants etc since it was utilized for species distribution prediction in 2004 and nowadays has universal applicability (Phillips *et al*., 2006; Wang *et al*., 2017). To expand the suitable growing region of the sweet cherry and avoid the investment risk of blind expansion, its potential distribution was modeled by using this model under current conditions based on these known coordinates and related environmental layers selected responsible for species distribution. The aims are: (1)to determine the potential growing areas of the species under the current condition; (2)to identify the most key variables responsible for the potential distributions, which will be helpful for the government at all levels to provide scientific guidance for carrying out reasonably regional planning and high-quality cultivation of this species in China.

## 2. MATERIALS AND METHODS

### 2.1 Species distribution samples

The main precise occurrence records from species specimen of *P. avium* were acquired from 4 free databases including: (1)the Chinese Virtual Herbarium (http://www.cvh.org.cn), (2)the National Specimen Information Infrastructure (http://www.nsii.org.cn/2017/home.php), (3)the Plant Photo Bank of China (http://ppbc.iplant.cn/), and (4)the Flora Reipublicae Popularis Sinicae (http://frps.iplant.cn/) etc. Moreover, some other occurrence records for this species were mainly collected from some scientific published article (Sun *et al*., 2021; Zhang *et al*., 2016; Nie *et al*., 2022; Cheng *et al*., 2023). In order to avoid some errors highly correlated with obvious misidentifications, some data that cannot obtain accurate geographical location were deleted, and some duplicates points data regarded as a error source owing to high spatial autocorrelation from the adjacent location (Jaryan *et al*., 2013) were removed too. Consequently, one occurrence point in each grid cell was ultimately retained. Finally, 422 occurrence points in total were eventually reserved and then the geographic coordinates of occurrence points were obtained based on the Gaode Pick Coordinate System (https://lbs.amap.com/tools/picker).or the Baidu Pick Coordinate System (http://api.map.baidu.com/lbsapi/getpoint/index.html). To be compatible with the software package Maxent, the coordinates of each point were kept in the csv format according to the species name, longitude and latitude in order.

### 2.2 Environmental data

The environmental variables including temperature, precipitation and topography factors etc. have the potential to determine species’ geographical distribution ranges (Wang *et al*., 2019; Qin *et al*., 2017). Firstly, we selected 19 bioclimatic variables with 30-arc-second (ca.1km^2^ at ground level) spatial resolution, under the current (i.e., in the period of 1970-2000) condition, downloaded from global Worldclim website (http://www.worldclim.org). These bioclimatic variables are equal to the average values from the years of 1970-2000, reflecting a combination of annual changes, seasonal characteristic, and extreme environmental conditions (Hijmans *et al*., 2005; Xu *et al*., 2014). Moreover, the altitude (DEM, Digital Elevation Model) data with the above-mentioned spatial resolution were also downloaded from the WorldClim website (http://www.worldclim.org) and were adopted to produce the slope and aspect data in ArcGIS 10.2. Secondly, the administrative boundary map of China (1: 400 million) was acquired from the national fundamental geographic information system (http://nfgis.nsdi.gov.cn/). Finally, these 22 environmental variables including 19 bioclimatic and 3 terrain variables were extracted by the boundary maps of China from the above-mentioned global raster data to ensure that all the research area have the same geographic bounds and cell size.

### 2.3 Variables selection

Due to the certain correlation between various environmental factors, if directly applied to the model, there may be overfitting phenomenon thought as error source (Jaryan *et al*., 2013; Jia *et al*., 2017). In order to eliminate this negative effects on model building and establish a model that has better performance with fewer variables, these 22 environmental variables were extracted from the corresponding layers associated with 422 documented occurrence points in ArcGIS 10.2 and then cross-correlations analysis (Pearson correlation coeffificient, *r*) have been performed. The decision to include or exclude one of each set of highly correlated variables was made based on their relative predictive power (Kumar *et al*., 2014). According to Pearson correlation coefficient (/r/ ≥ 0.8) in SPSS 21.0 and taking into consideration their relative predictive power (i.e., the importance of each environmental variable to predictor contributions from analysis of predicted results), only one variable with higher percent contribution from each set of highly cross-correlated variables (/r/≥0.8) was kept in further analyses while the other variable was excluded because of its lower predictive power. Otherwise, all environmental factors were retained. Finally, a total of remaining 11 variables were kept and utilized for model building (Table 1). Meanwhile, to meet the needs of this model, all these remaining variables were converted to asc formats.

**Table 1.** The environmental factors used in this study, their contribution and permutation importance under current environmental condition.

| Code | Description | Percent contribution | Permutation importance | Code | Description | Percent contribution | Permutation importance |
| --- | --- | --- | --- | --- | --- | --- | --- |
| bio06 | Min temperature of coldest month | 50.9 | 56.3 | Bio14 | Precipitation of driest month | 2.5 | 3.4 |
| bio12 | **Annual precipitation** | **22.9** | **12.5** | bio03 | Isothermality | 0.8 | 1.7 |
| bio10 | Mean temperature of warmest quarter | **9.5** | **13.4** | Alt | alt30s1 | 0.4 | 2.2 |
| Slo | slope | 5.5 | 4.4 | Asp | aspect | 0.2 | 0.2 |
| bio15 | Precipitation Seasonality | 4.3 | 2.1 | bio02 | Mean diurnal range | 0.1 | 1.0 |
| bio19 | Precipitation of driest month | **2.7** | 2.9 |  |  |  |  |

Note: the highlighted variables, selected through their contribution and permutation importance, were five main influencing factors.

### 2.4 Modeling procedure

The potential geographical distributions of *P. avium* were modeled by the Maxent model (Maximum entropy model)(Version 3.4.1) downloaded freely for scientific research from http://www.cs.princeton.edu/∽schapire/maxent/, which was widely utilized to quantify the impact of climate and other environmental changes on species distributions and get good performances (Qin *et al*., 2017; Wang *et al*., 2017). In the process of model building, 422 known occurrence data of *P. avium* and 11 environment variables can be directly imported into the corresponding module of Maxent model, wherein, the training data were 75% of all the occurrence data selected at random, and the test data were the remaining 25% (Wang *et al*., 2010; Qin *et al*., 2017). Meanwhile, a jackknife test and response curves modules were checked in the model interface to calculate the habitat suitability curves and estimate the importance of each environmental variable that affect this species distributions (Kumar *et al*., 2014). The maximum number of background points was set up to 10,000. The other settings were the same as described in some scientific published article (Li *et al*., 2021). In order to guarantee the steadiness of prediction results, the model was run 10 replicates by cross-validation and the average habitat suitability was regarded as the final result in logistic format and asc types (Khanum *et al*., 2013; Zhang *et al*., 2022). Finally, this result was transformed into raster format and the cell value of prediction results ranges from 0 representing the lowest habitat quality for that species, to 1 representing the highest habitat quality. The Maximum Youden Index (Maximum training sensitivity plus specificity logistic threshold) and the TPT equilibrium threshold (Balance training omission, predicted area and threshold value) were usually used to be the cutoff points, which are advantage over other threshold value (Jimenez-Valverde and Lobo, 2007; Yang *et al*., 2020). Finally, based on the above cutoff points, the continuous habitat areas for *P. avium* were divided into 3 categories, such as suitable, medium and unsuitable area.

### 2.5 Model performance and influencing factors

The area under the receiver operating characteristic curve (the AUC) is usually used to assess model’s goodness-of-fit and nowadays the AUC value was widely regarded as an excellent index in evaluating model performance (Qin *et al*., 2017; Wang *et al*., 2021). The software package Maxent (Version 3.4.1) used in our study makes use of the AUC to assess its performance. The AUC value varies from 0.5, implying that the prediction performance was not better than that of the random model or the lowest predictive ability, to 1.0 showing the best performance or the highest predictive ability (Xu *et al*., 2014; Yang *et al*., 2020). Model performance may be classified as excellent (0.9–1), good (0.8–0.9) or ordinary (0.7–0.8), respectively (Yi *et al*., 2016). According to our regulations, Models of this species with values above 0.8 were regarded useful in our study whereas AUC values less than 0.8 were not considered in subsequent research. In general, the larger the AUC is, the better the model performance is (Yi *et al*., 2016). By using the software’s built-in jackknife test, based on the predictor contributions, permutation importance and regularized training gain, we can be able to assess the relative influence of individual predictors to the species’ habitat suitability (Dong *et al*., 2019; Zhang *et al*., 2022; Wang *et al*., 2021).. Furthermore, the response curve generated automatically by Maxent model is used to show the quantitative relations between the environmental variables and the logistic probability of occurrence (Hu *et al*., 2015; Li *et al*., 2021).

## 3. RESULT

### 3.1 Modeling evaluation

The geographic distribution map was generated by Maxent model based on 422 known occurrence data of the *P. avium* and 11 environment variables selected. Results showed that the AUCs of modeling and testing are near 1.0 (0.9449±0.001 for training and 0.9301±0.0112 for testing) under the current conditions (Table 2), suggesting that Maxent model performed excellent in predicting the geographic distribution area for the *P. avium*.

**Table 2.** The AUC, Maximum Youden Index and TPT equilibrium threshold generated in 10 replicates.

| No. | AUC of model building | AUC of Model testing | Maximum Youden Index | TPT equilibrium threshold |
| --- | --- | --- | --- | --- |
| 1 | 0.9456 | 0.9264 | 0.2483 | 0.0427 |
| 2 | 0.9465 | 0.9124 | 0.2317 | 0.0274 |
| 3 | 0.9443 | 0.9434 | 0.2181 | 0.0402 |
| 4 | 0.9441 | 0.944 | 0.182 | 0.0477 |
| 5 | 0.9441 | 0.9361 | 0.2115 | 0.0499 |
| 6 | 0.9435 | 0.9429 | 0.1892 | 0.0554 |
| 7 | 0.9458 | 0.9168 | 0.2374 | 0.0502 |
| 8 | 0.9461 | 0.9182 | 0.2126 | 0.0536 |
| 9 | 0.9441 | 0.9365 | 0.1755 | 0.0377 |
| 10 | 0.9449 | 0.9244 | 0.1901 | 0.0427 |
| Average | 0.9449 | 0.9301 | 0.2096 | 0.0447 |

Note: Maximum Youden Index (Maximum training sensitivity plus specificity logistic threshold) and TPT equilibrium threshold (Balance training omission, predicted area and threshold value) in analysis of predicted results.

### 3.2 Dominant environmental factor restricting the distribution of P. avium in China

Among the 11 environmental variables, the min temperature of coldest month (bio06), the annual precipitation (bio12) and the mean temperature of warmest quarter (bio10) were the strongest predictors associated with the potential distribution of *P. avium* relative to other variables according to percent contribution (Table 1). The min temperature of coldest month (bio06) made the largest contribution (50.9%), indicating that this variable significantly has an effect on the distribution of *P. avium* under the current condition, followed by the annual precipitation (bio12) and the mean temperature of warmest quarter (bio10), with 22.9% and 9.5% respectively. The cumulative contributions of these factors reached values as high as 83.3%. Meanwhile, based on permutation importance (Table 1), the min temperature of coldest month (bio06) with 56.3%, had the highest score (Table 1), followed with the annual precipitation (bio12) and the mean temperature of warmest quarter (bio10), with 12.5% and 13.4% respectively. The cumulative permutation importance of these three parameters was equal to 82.2%. In addition, based on the software’s built-in jackknife test (Fig. 1), the min temperature of coldest month (bio06), the annual precipitation (bio12) and the mean temperature of warmest quarter (bio10) had the highest predictive power (highest regularized training gain)on the geospatial distribution of the *P. avium*. Taken together, three dominant environmental variables identified were the min temperature of coldest month (bio06), the annual precipitation (bio12) and the mean temperature of warmest quarter (bio10).

**Fig. 1.**
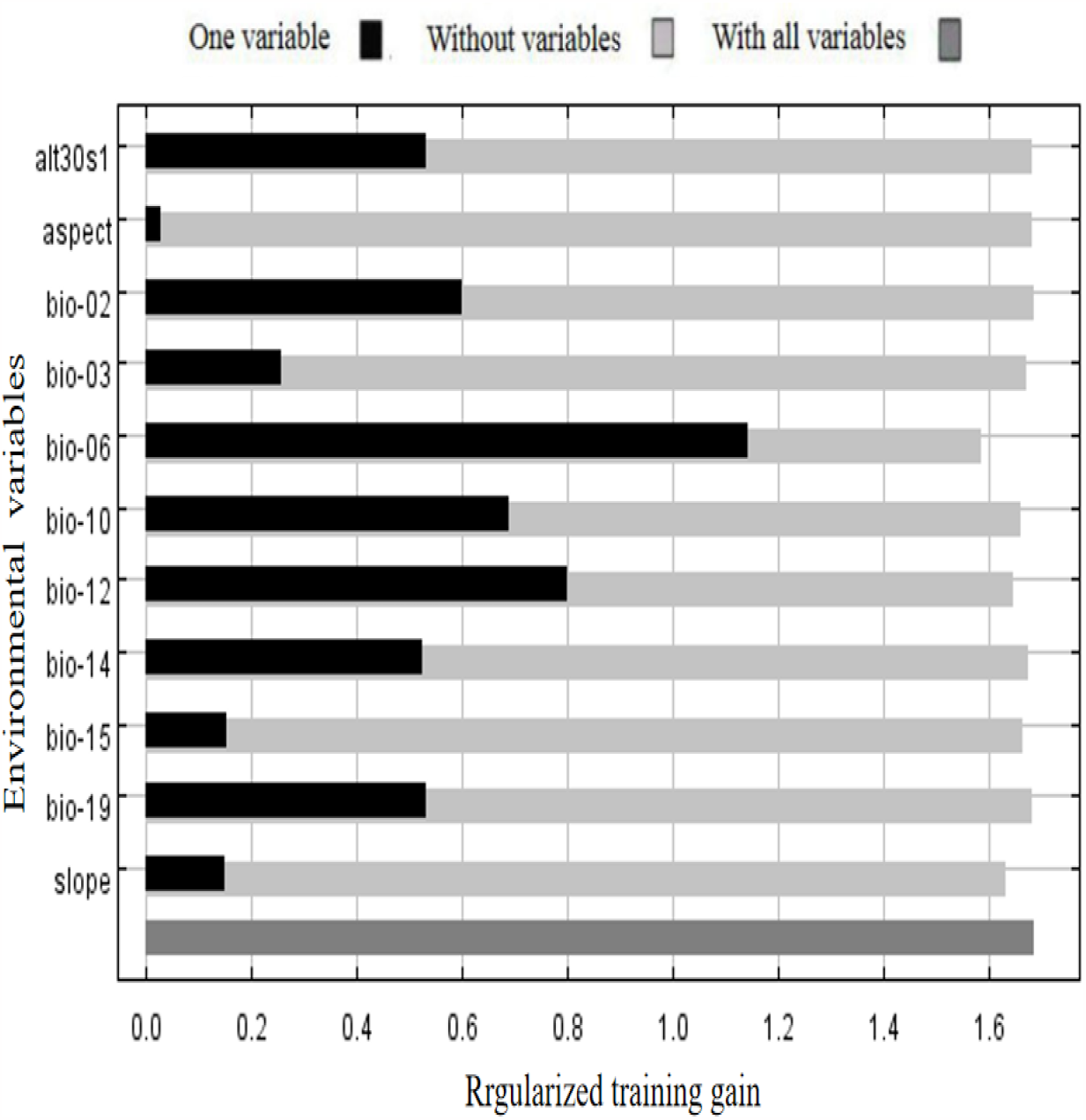
Jackknife test for assessing relative importance of different environmental variables v to the geospatial distribution of *P. avium* under the current condition.

### 3.3 *Environmental characteristics of* the *P. avium in China*

The area with a fitness level greater than 0.2096 (the Maximum Youden Index) is the simulated high fitness area of the *P. avium*, which matched closely with the actual distribution area (the documented occurrence of the *P. avium*) (Fig. 3), implying that the above-mentioned threshold value is applicable. Therefore, in this study, threshold of all environmental factors was used to show the characteristics of the distribution area for this species in China when the probability of existence was greater than 0.2096. Furthermore, to clarify the environmental characteristics of this species’ distribution and obviate interrelated interference among environmental factors, the above 3 key factors were alone imported into the MaxEnt model. Finally, individual response curves showed the quantitative relationship between each environmental factor and the logistic probability of presence. As shown in Fig. 2, the min temperature of coldest month (bio06) ranging from -14.5 to 4.5°C, the most highest point is -6.5°C; the annual precipitation (bio12) from 500 to 1200 mm; the mean temperature of warmest quarter (bio10) from 21.0 to 28.0°C.

**Fig. 2.**
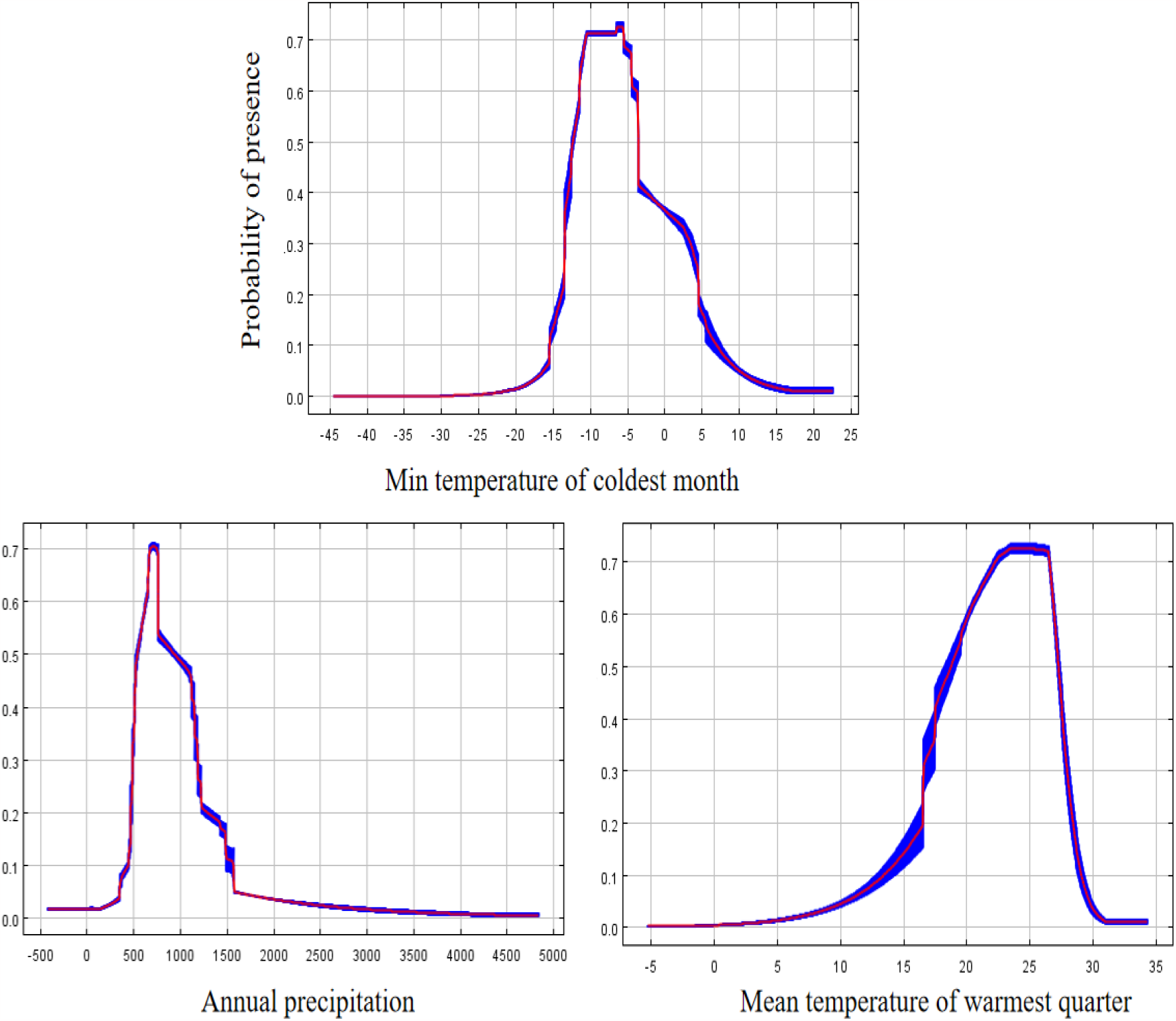
Response curves of Maxent jackknife method and main environment variables.

**Fig. 3.**
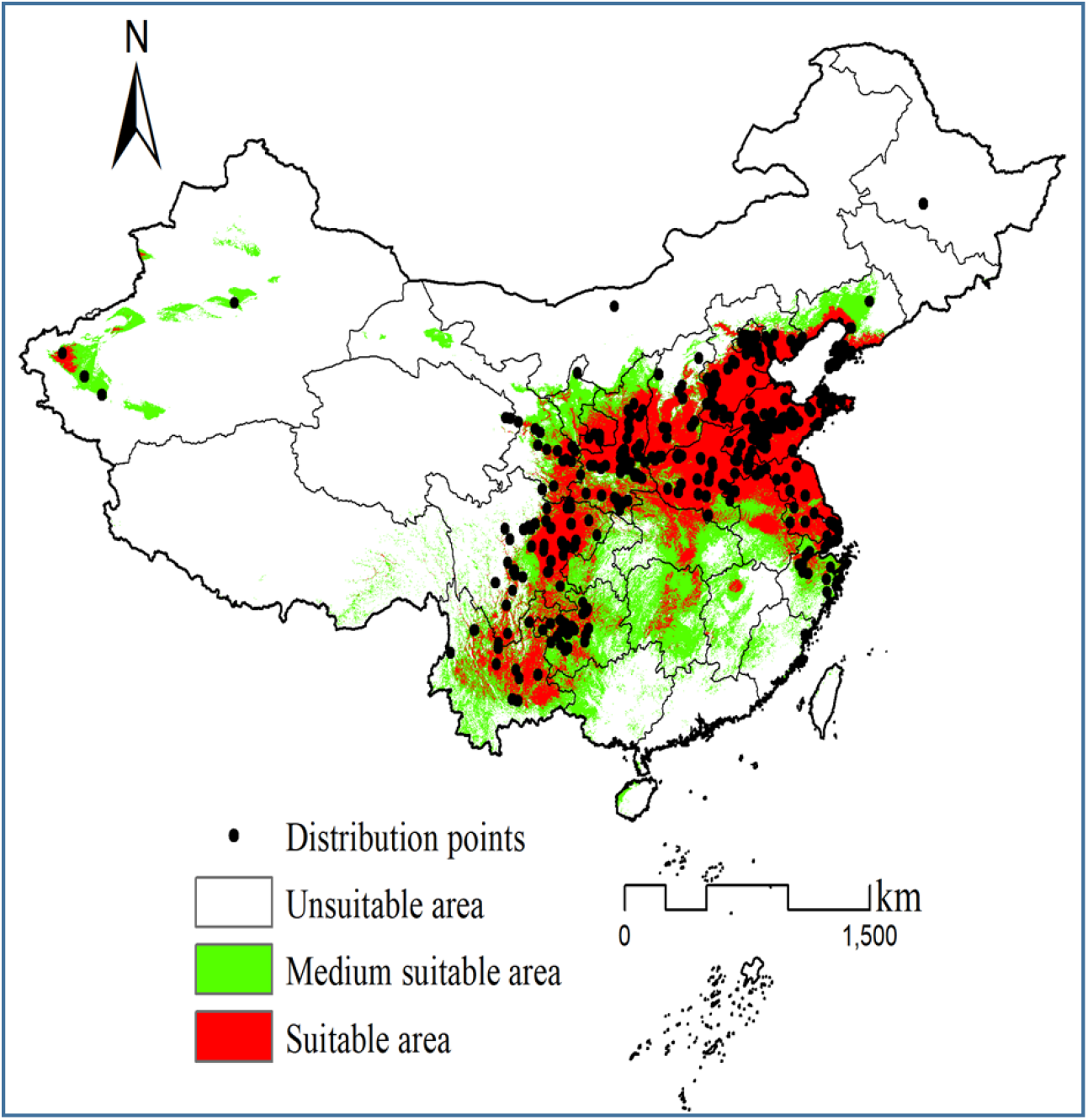
The potential geospatial distribution of *P. avium* in the periods of 1970-2000 in China.

#### Predicting potential distribution

Based on the Maximum Youden Index (the average value of 0.2096 in 10 replicates) and TPT equilibrium threshold value (the average value of 0.0447 in 10 replicates) as a threshold value under the current condition (Table1), the final distribution map was reclassified into three categories: unsuitable area (<0.0477), medium suitable area (0.0447∽0.2096) and suitable area (> 0.2096). The suitable region of the *P. avium* in the study region mainly is concentrated on six areas including southwest China (eastern Sichuan, northeast and and main urban areas of Chongqing, mid-western Guizhou and mid-northern Yunnan), northwest China (mid-southern Shaanxi, southern Ningxia mid-southern and eastern Gansu), northeast China (Coastal region of Liaoning), central China (most of Henan, mid-northern Hubei and central Hunan), north China (Beijing, Tianjing, mid-southern Shanxi), east China (Shanghai, Jiangsu, Shandong, central Zhejiang, central and northern Anhui and eastern Jiangxi) and south China (western Guangxi). The medium suitable area lies in southwest China (Chongqing city, Guizhou, eastern Guizhou, western and southern Yunnan and sporadic areas in eastern Sichuan), northwest China (northern Shaanxi, mid-southern Gansu and Ningxia), northeast China (mid-western of Liaoning), central China (sporadic areas in Henan, western and eastern Hubei and most of Hunan), north China (mid-northern Shanxi and eastern Hebei), east China (most of Zhejiang, central and southern Anhui, and most of Jiangxi except for eastern region) and south China (western Guangxi and northwest Hainan) (Fig. 3). Except for the above regions, other regions in the research region are not suitable for the growth of *P. avium*. This part of the region mainly includes most of Tibet, western of Ningxia, Qinghai, Xinjiang, Inner Mongolia, Heilongjiang, Jilin, Guangdong, Hainan and Taiwan of China. According to the statistical analysis after the projection conversion (Fig. 3) (Asia_North_Albers_Equal_Area_Conic), the percentages of suitable, medium and unsuitable growing regions of *P. avium* were 6.81%, 13.51% and 79.68% in China, respectively. However, for various regions in China (Table 3), the potential suitable area of *P. avium* is different under the current condition. For example, the percentages of the suitable areas in 13 provinces or cities including Sichuan, Guizhou, Yunnan, Shaanxi, Beijing, Tianjing, Shanxi, Hebei, Henan, Hubei, Shanghai, Jiangsu and Shandong reach over 20% respectively, while those in Chongqing, Gansu, Ningxia, Hunan, Zhejiang, Anhui, Jiangxi, Hainan and Guangxi provinces or cities are less than 20%, however, these regions have some moderate adaptation areas. Moreover, compared with their actual distribution, Hubei, Hunan and Anhui etc also have some potentially suitable areas. Among them, those in Shaanxi, Beijing, Tianjing, Shanxi, Hebei, Henan, Shanghai, Jiangsu and Shandong etc reach over 48.66% while Sichuan, Guizhou,Yunnan, Gansu, Liaoning, Hubei, Zhejiang and Guangxi etc ranged from 17.10∽29.44% respectively (Table. 3), indicting these 9 provinces or cities are main regions for its current development and utilization, followed by Sichuan, Guizhou,Yunnan, Gansu, Liaoning, Hubei, Zhejiang and Guangxi etc. The medium suitable area mainly exists in Chongqing, Guizhou,Yunnan, Ningxia, Liaoning, Hubei, Hunan and Zhejiang etc based on medium percentage of predicted area.

**Table 3.**
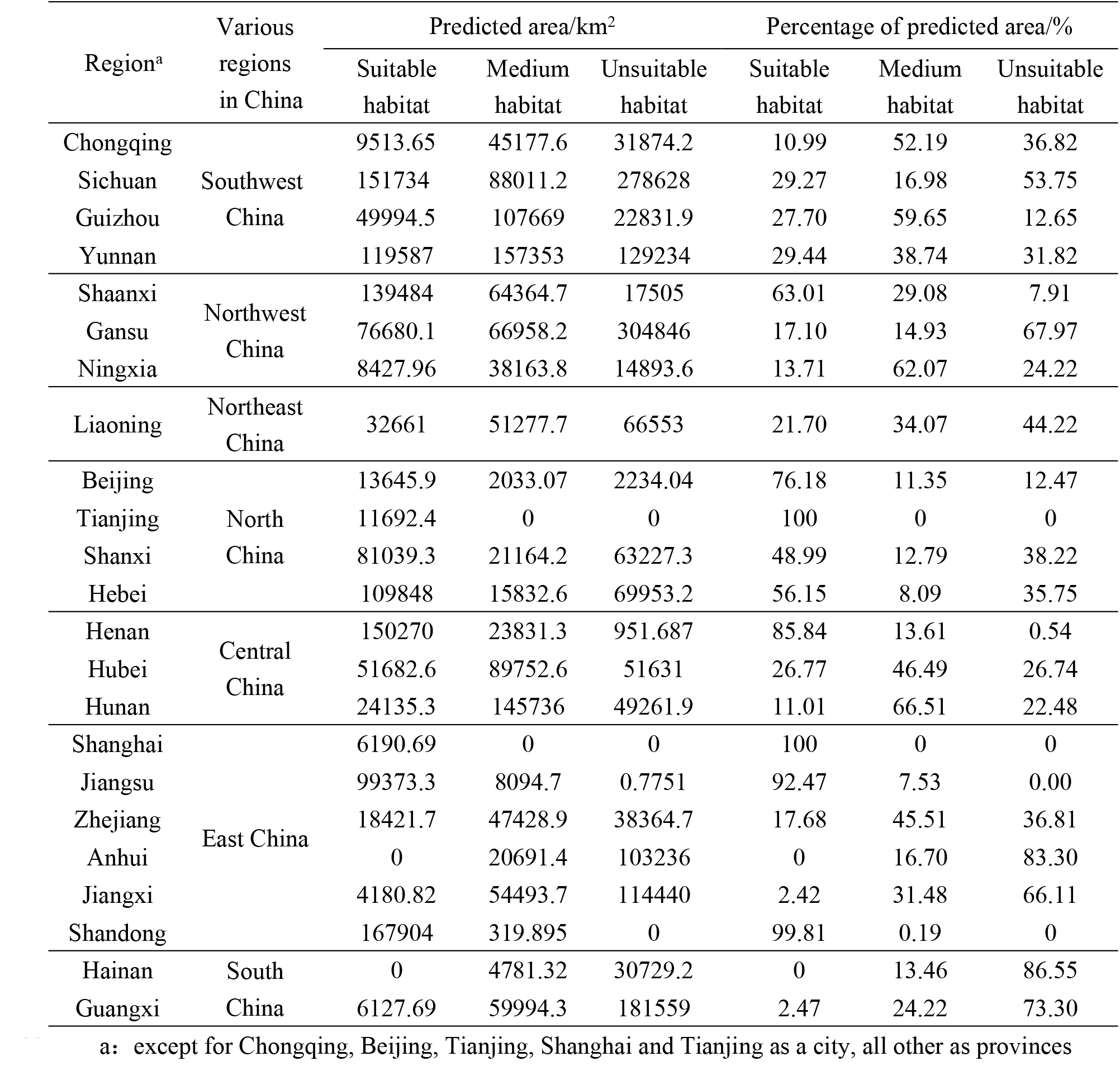
Area and percentage of habitat distribution for *P. avium* in different provinces or cities under the periods of 1970-2000.

## 4 Discussion

The sweet cherry with rich bioactive substances such as vitamin C and polyphenols etc. is known as “vitamin pills” by experts (Wang, 2014; Commisso *et al*., 2017). It indicated that it has still high medicinal value (Wu *et al*., 2018). If the human body lacks the above elements, the corresponding diseases will occur, such as iron deficiency causing anemia, vitamin A deficiency causing dry skin, and vitamin C deficiency causing gum bleeding. Meanwhile, they are a health fruit that prolongs people’ life, especially with significant beauty effects, so they are very popular among women and present a strong sales momentum (Wang, 2014). However, for a long time, due to the intolerance of storage and transportation, their cultivation ranges have been very narrow, resulting in slow expansion speed and limited yield, which cannot meet market needs. To meet the increasing demand for the sweet cherry, their cultivation ranges and production scale in China are continuously expanding with the improvement of cultivation technology and the rapid development of modern storage and transportation industry. However, so far, very little is aware of the potential suitable areas of the sweet cherry, which results in the investment risks mainly evoked by blindly expanding its introduction and growing area. Fortunately, many SDMs have been widely adopted to model the potential distributions of many species and get good prediction results (Wang *et al*., 2010; Yi *et al*., 2016). Wherein, the predictive ability of Maxent model was always stable and reliable, and it outperformed other SDMs (Elith *et al*., 2006). In this study, its potential distributions were modeled under current conditions based on only 422 precise coordinates of species occurrences (In practice, it is very difficult to obtain absence data) together with 11 environmental layers, their prediction performances are very excellent, with AUCs of greater than 0.9 for modeling and testing. Furthermore, based on above-mentioned predicting outcomes (Fig. 3), the suitable area matched closely with the actual distribution of the sweet cherry, indicating that their predicted results were precise and the above threshold value under the current condition was reliable. It is reported that in China there are two advantageous cultivation areas for the sweet cherry. One is the Bohai Bay Rim area, which is dominated by Shandong (Yantai and Taian) Liaoning (Dalian), Beijing and Hebei (Qinhuangdao), the other is the area along the Longhai railway, dominated by Henan (Zhengzhou), Shaanxi (Xi’an), and Gansu (Tianshui) (Zhang *et al*., 2017). These results are substantially similar to our predicted results based on this model. However, the model also identified some potentially suitable areas for the sweet cherry in China, which indicates that the sweet cherry has not yet reached its full potential range (Fig. 3). As shown in Fig. 3 and Table 2, in southwest China, the percentages of the suitable growing areas in Sichuan, Guizhou and Yunnan provinces respectively reach over 20% while it only is 10.99% in Chongqing city; In northwest China, those in Gansu and Ningxia provinces are less than 20% while it reach 63.01% in Shaanxi; In northeast China, it is only 21.70% in Liaoning; In north China, those in Beijing, Tianjing, Shanxi and Hebei respectively reach over 20%; In central China, those in Henan and Hubei provinces will reach over 20% while it reach 11.01% in Hunan; In east China, those in Shanghai, Jiangsu and Shandong respectively reach over 20% while those in Zhejiang, Anhui and Jiangxi are less than 20%; In south China, those in Hainan and Guangxi are less than 20%. Through comprehensive analysis, those in Shaanxi, Beijing, Tianjing, Shanxi, Hebei, Henan, Shanghai, Jiangsu and Shandong etc reach over 48.66% while Sichuan, Guizhou,Yunnan, Gansu, Liaoning, Hubei, Zhejiang and Guangxi etc ranged from 17.10∽29.44% respectively (Table 2), indicting these 9 provinces or cities are main regions for its current development and utilization, followed by Sichuan, Guizhou,Yunnan, Gansu, Liaoning, Hubei, Zhejiang and Guangxi etc. In addition, the percentages of the medium growing areas in Chongqing, Guizhou, Yunnan, Shaanxi, Ningxia, Liaoning, Hubei, Hunan, Zhejiang, Jiangxi, and Guangxi provinces or cities respectively ranged from 24.22∽62.07% (Table 2), indicting these provinces or cities may introduce some special products for the local development.

The potential distribution area of species was strongly correlated with environmental variables including temperature, rainfall, terrain and so on (Jia *et al*., 2017; Zhang *et al*., 2022). Among environmental variables, climate (temperature and rainfall) is among the most critical factors limiting the potential distribution of species(Zhang *et al*., 2022). Based on the predictor contributions, permutation importance and regularized training gain, these dominant environmental variables identified were temperature (the min temperature of coldest month (bio06) and the mean temperature of warmest quarter (bio10)) and rainfall (the annual precipitation (bio12)) variables, wherein the min temperature of coldest month (bio06) made the greatest contributions to the distribution model for the *P. avium*, indicating that thermal factors are the dominant climate factor affecting distribution of the sweet cherry in China, followed by water factors during the growing season. It was reported that the critical low temperature during the winter dormancy period of sweet cherry should not be lower than -20°C, which is considered the critical temperature for the occurrence of winter damage in large cherries (Yao *et al*., 2009; Yan *et al*., 2013) . On the contrary, if below this critical temperature, it is difficult to safely overwinter, such as its main trunk and branches are prone to frost cracking, gummosis and flower buds are also susceptible to frost damage; at -25 °C, it causes the entire cherry tree to die, which limits the northward development of sweet cherry. As for Yunnan and Guizhou provinces etc., the main factor restricting the cultivation of sweet cherry is the insufficient cold demand in winter, so the fruit tree often blooms but does not bear fruit (Zhang *et al*., 2016). If the temperature is too high in summer, cherry trees exhibit excessive vigor, branches stop growing later, canopy closure is elevated, so fruit quality is poor, resulting in loss of commercial value. Most seriously, high temperatures in summer can cause abnormal differentiation of flower buds in the early stages to produce a large number of deformed flowers, such as the twin-room flower, it will form a “twin fruit” in the coming year. Further it is proved that the sweet cherry is suitable to live in the min temperature of coldest month (bio06) from -14.5 to 4.5°C,the most highest point is -6.5°C, which can ensure maximum flowering and fruiting; and the mean temperature of warmest quarter (bio10) from 21.0 to 28.0°C. In addition, a lots of sweet cherries are suitable for growing in areas with annual precipitation of 600∽800 mm through past research (Zhang *et al*., 2016; Cheng *et al*., 2023) . This was highly consistent with with our research results, this was to say, the annual precipitation (bio12) ranged from 500 to 1200 mm.

## Conclusion

(1)The suitable area of the *P. avium* in China mostly is located in southwest China (eastern Sichuan, northeast and and main urban areas of Chongqing, mid-western Guizhou and mid-northern Yunnan), northwest China (mid-southern Shaanxi, southern Ningxia mid-southern and eastern Gansu), northeast China (Coastal region of Liaoning), central China (most of Henan, mid-northern Hubei and central Hunan), north China (Beijing, Tianjing, mid-southern Shanxi), east China (Shanghai, Jiangsu, Shandong, central Zhejiang, central and northern Anhui and eastern Jiangxi) and south China (western Guangxi). (2)Based on the percentages of its suitable growing regions, these 9 provinces or cities (Shaanxi, Beijing, Tianjing, Shanxi, Hebei, Henan, Shanghai, Jiangsu and Shandong) are main regions for its current development and utilization, followed by Sichuan, Guizhou,Yunnan, Gansu, Liaoning, Hubei, Zhejiang and Guangxi etc. (3) The key factors selected were the min temperature of coldest month (bio06) ranged from -14.5 to 4.5°C, the most highest point is -6.5°C; the annual precipitation (bio12) from 500 to 1200 mm; the mean temperature of warmest quarter (bio10) from 21.0 to 28.0°C. And, thermal factors are the dominant climate factor affecting distribution of the sweet cherry in China, followed by water factors during the growing season.

## Acknowledgements

The reviewers sincerely were thank for the important comments on our paper that heightened my paper level. We especially thank Shaobin Gao for assistance in data collection and organization. The research was financially aided by the National Natural Science Foundation of China (31870515), Rescue and Protection Projects for Rare and Endangered Wild Fauna and Flora in Chongqing Municipality (2023-1) and Chongqing Natural Science Foundation(CSTB2023NSCQ-MSX0591).

## Authors’ contribution

HL, XP, PJ, and LX designed the research.

HL implemented the research.

HL wrote the paper.

## Conflict of interest disclosure

All authors declare that there is no confrontation of benefit.

## Notes

### Competing Interest Statement

The authors have declared no competing interest.

